# A machine learning approach to identify key Epigenetic Transcripts for Ageing research in human blood (Epitage)

**DOI:** 10.64898/2026.02.09.704870

**Authors:** Thiago Benazzi Maia, Ulrich Pfeffer

## Abstract

DNA methylation is an established biomarker of human ageing, and analysing CpGs grouped by transcript as functional units may reveal new insights into the processes of ageing. In this study, we analyzed the **GSE87571** dataset (714 samples from 14–94 years) to assess the relationship between transcript-level methylation profiles and chronological age in human blood. This approach led to the creation of **Epitage**, a curated set of 48 transcripts from 13 genes identified through machine learning as having methylation profiles that strongly correlate with age (R^2^ ≥ 0.8). This analysis highlighted transcripts from the genes **KCNS1, SPTBN4**, and **VTRNA1-2**, which have been only rarely mentioned as age-related methylation markers in humans, suggesting them as underexplored candidates for future investigation. In addition, the list includes genes already implicated in aging or related pathways, such as **ELOVL2, FHL2, KLF14, TRIM59, MIR29B2CHG, CALB1, OBSCN, PRRT1, OTUD7A, and SYNGR3**. To validate models efficiently while ensuring reproducibility, we developed **ugPlot**, an open-source R package with a graphical user interface (GUI) that automates routine steps for training and testing hundreds of machine-learning models. The tool also streamlines dataset import and manipulation, reducing human error and generating publication-ready plots. Epitage thus provides a focused and accessible starting point for experimental and translational studies into the roles of DNA methylation and transcript regulation in human ageing.

## 1 Introduction

Ageing manifests at the cellular level through deregulation of signaling pathways [43, 45], cumulative damage and epigenetic drift [23]. Although the accumulation of such alterations has been frequently reported, a fundamental question remains unresolved: are DNA methylation changes a cause of ageing related functional decline, merely a consequence of underlying cellular deterioration or perhaps an adaptive response to cellular stress? Distinguishing correlation from causation is critical. If methylation shifts are merely markers, they may be useful for diagnostics, but if they are drivers, they become viable therapeutic targets [31].

This uncertainty about causality highlights the need for functional experiments. If age-associated transcripts, identified through methylation patterns, are shown to directly influence ageing phenotypes, then restoring their expression could have therapeutic relevance. Unlike genetic mutations, which are typically permanent and relatively rare, DNA methylation is of particular interest due to its plasticity, stability, and heritability during cell division [4].

Accurately assessing ageing requires distinguishing between chronological age, defined as the time since birth, and biological age, which reflects the extent to which an individual shows signs of ageing compared to others of the same chronological age [22]. This distinction is not merely conceptual, but could have clinical relevance: several studies have suggested that accelerated epigenetic ageing; as measured by deviations in DNA methylation patterns from expected chronological values [22, 46]; may be associated with increased risk of age-related conditions, including cardiovascular disease [34], cancer [34], neurodegenerative disorders [27], type 2 diabetes [7], and frailty [35]. Ageing biomarkers may therefore have potential not only as indicators of time but also as tools for risk assessment in age-related diseases.

While whole genome sequencing can detect somatic mutations that accumulate in cells over time [13], epigenetic changes, particularly DNA methylation, are often described as more dynamic and potentially influenced by environmental and lifestyle factors [14]. CpG methylation plays a role in regulating chromatin structure and gene expression [5]. Methylation patterns vary between cell types, yet tend to remain relatively consistent within a given cell population [19], which makes them suitable for use as biomarkers in ageing research.

When discussing methylation, the term CpG is often referred to, which occurs in DNA when a Cytosine (C) is followed by a Guanine (G) on the same strand and linked by a phosphate group (p) [29]. In methylation, a methyl group (CH_3_) is added to the 5th position of the cytosine ring in DNA, forming 5-methylcytosine. Generally, the closer it is to the promoter, the greater the chance of influencing gene expression; that is, methylation works to block gene expression [41]. This study highlights that methylation within the gene body, beyond the promoter region, can also be functionally relevant and should be carefully considered.

Among emerging biomarkers, DNA methylation at CpG sites DNA has been shown to be both robust and reproducible. Epigenetic clocks such as Horvath’s Clock [22], Hannum’s model [20], PhenoAge [28], and GrimAge [32] use methylation patterns to estimate biological age. The difference between epigenetic and chronological age is often viewed as a potential indicator of increased risk for mortality and age-related diseases [28]; [32]; [9]. Yet, the causal nature of this association remains unclear, positioning DNA methylation as a promising, though still not fully understood, marker of biological health.

Ageing clocks are powerful tools for estimating biological age and have shown strong performance across multiple datasets. They typically rely on hundreds of CpG sites selected for predictive accuracy.

In addition to CpG-centric models, this study explores the potential of focusing on transcript-level methylation to gain biologically meaningful insights into ageing. We aim to identify gene transcripts whose methylation profiles are strongly and reproducibly associated with chronological age. This perspective can also support experimental approaches, such as reintroducing underexpressed transcripts into cells, to investigate whether restoring their activity affects ageing-related processes. This strategy may offer a safer and more targeted alternative than attempting to modify DNA methylation directly. Given the complexity of gene regulation, a curated list of age-associated transcripts can serve as a practical resource for future research and therapeutic development.

After extensive selection, we chose the publicly available dataset GSE87571 from the Gene Expression Omnibus (GEO), [1]. This dataset consists of 732 whole-blood samples, aged 14 to 94 years, with high-quality curation and metadata sufficient for the purpose of this research. The samples were derived from peripheral blood leukocytes and analyzed using the Illumina Infinium 450K Human Methylation BeadChip, a widely used platform that measures methylation at approximately 450,000 CpG sites with high reproducibility. Methylation levels are reported as *β*-values, representing the proportion of methylated alleles at each site [39]. Additionally, the dataset includes raw IDAT files (RED and GRN channels), allowing us to reprocess the data using modern quality control pipelines, such as those implemented in the **SeSAMe** [48] R package, ensuring more accurate estimates of biological age and robust filtering of inconsistent samples.

To efficiently manage large volumes of CpG data, we used MySQL as a database backend. After importing and indexing the dataset, query and analysis times were significantly reduced. This made it easier to extract CpGs associated with each transcript and organize the data for model testing. The database dump is available in the paper repository for reproducibility and further exploration.

The initial filtering of age-correlated transcripts was performed using R scripts, but as the analysis scaled up, a more robust solution became necessary. To address this, we developed **ugPlot** (https://github.com/a00s/ugplot), an R package available in the project repository, designed to simplify and accelerate model testing. ugPlot integrates over 200 machine learning models from the **Caret** [25] package, applies 10-fold cross-validation, and supports automated, seed-based reproducibility. It also includes a **Shiny**-based [8] visual interface and allows batch testing across multiple random seeds. This tool was essential for extracting reliable transcript– CpG associations without the burden of manually handling multiple files, custom code, and potential human errors.

Taking into account that some transcripts correspond to isoforms of the same gene [42], we grouped them accordingly to produce a curated list of 48 transcripts, referred to as Epitage. Each transcript is associated with a set of CpG methylation profiles that exhibit correlations (R^2^ ≥ 0.8) with chronological age in blood-derived DNA. When analyzed using the R package ugPlot, these entries offer an interpretable resource that may support investigations into agerelated epigenetic changes. The underlying hypothesis is that having such a list could guide future studies aiming to introduce synthetic versions of downregulated, age-associated transcripts into cells. This strategy might represent a relatively safer and more controllable alternative, at least provisionally, compared to attempting direct modification of DNA methylation patterns within cells.

In conclusion, this study presents a resource for investigating the epigenetic regulation of ageing at the transcript level. The combination of ugPlot with the Epitage list provides a starting point for exploring how specific methylation profiles may relate to age-associated changes in gene regulation. Rather than offering a definitive catalog, Epitage should be viewed as an evolving reference, one that can be refined as new transcripts are implicated in ageing and as more comprehensive or higher-resolution methylation data become available through advances in technology.

## 2 Materials and Methods

### 2.1 Overview of the process

The workflow was divided into three main stages, as described in Figure 1, to ensure methodological consistency and robust validation.

**Figure 1.**
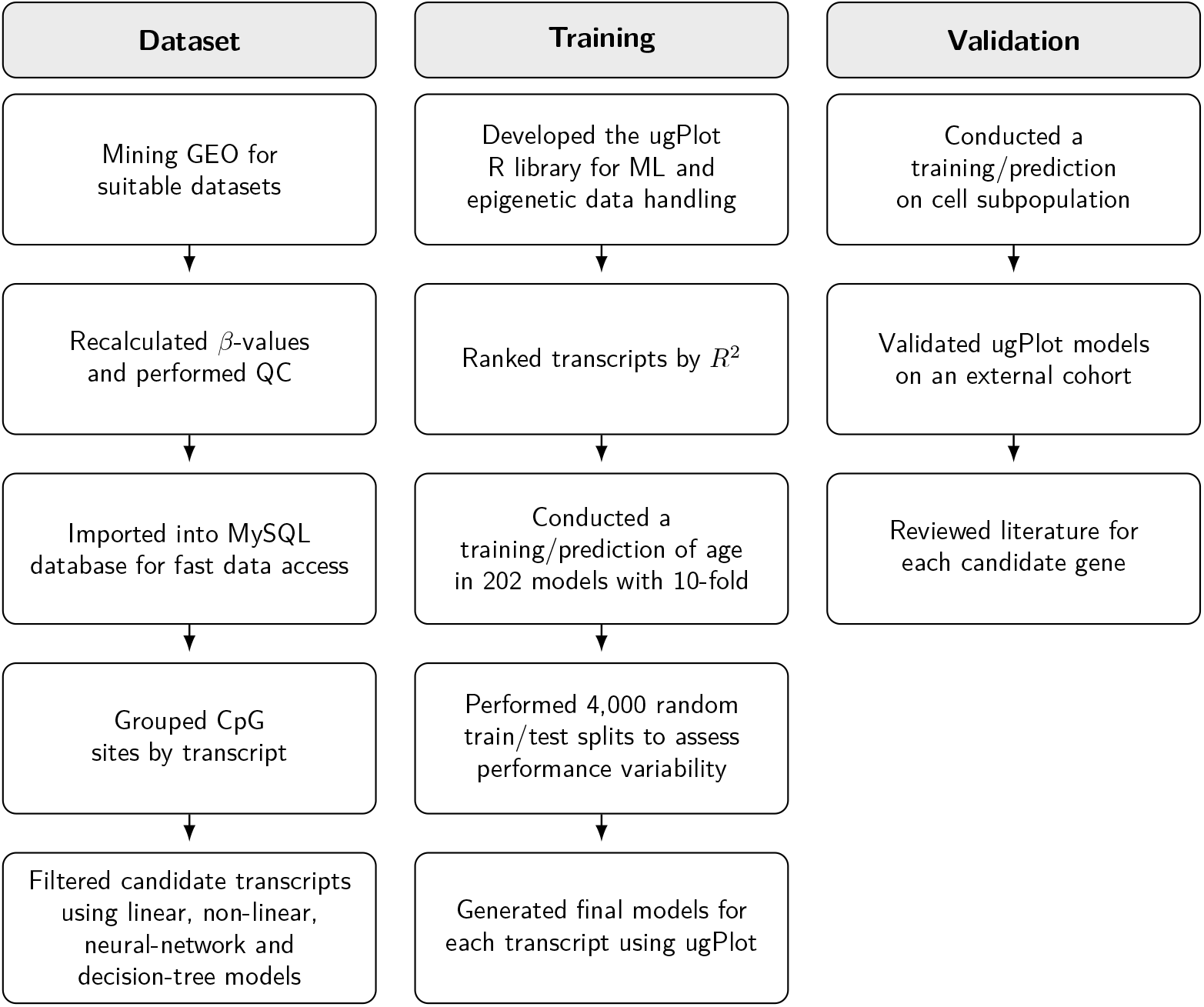
Overview of the process by which this study was conducted, from dataset preprocessing to model training and external validation using the **ugPlot** R package.

### 2.2 Dataset

For this study, we used the publicly available GEO dataset GSE87571 [1], “The relative contribution of DNA methylation and genetic variants on protein biomarkers for human diseases”), which provides DNA methylation profiles generated with the Illumina Infinium HumanMethylation450 BeadChip (450K array), covering more than 450,000 CpG sites across the genome. The dataset consists of 732 whole-blood samples, aged 14 to 94 years and documented sex. All raw data from GSE87571 were reprocessed with quality control and *β*-value recalculation as detailed in the Methods section to ensure consistency for downstream analyses. This wide age range and detailed metadata make GSE87571 particularly well-suited for exploring age-associated methylation patterns in human blood. Ideally, we would have excluded samples from the same individual collected at different ages to avoid repeated measures, but this was not possible due to insufficient metadata. To mitigate this limitation, we applied additional validation methods to ensure our results were not solely driven by model artifacts.

For external validation, we used data from the study “Exposure to polybrominated biphenyl (PBB) associates with genome-wide DNA methylation differences in peripheral blood” [10]; GEO accession GSE116339; which includes 673 peripheral blood samples from 658 participants, aged 23 to 88 years. Although this dataset consists of individuals exposed to PBB, which could potentially introduce some bias or imprecision in the results, the high quality of the data made it a valuable resource for validating our findings in an independent cohort. Unlike the main study dataset, for GSE116339 we did not perform any additional *β*-value recalculation or quality control; the data were analyzed exactly as published in order to test the reproducibility and robustness of our results.

### 2.3 Preprocessing and Quality Control

Raw intensity data (.idat files) from the 450K array (GEO GSE87571) were processed in R using the SeSAMe package [48]. *β*-values, representing the fraction of methylated DNA at each CpG, were calculated after performing all essential preprocessing steps: background correction (to reduce technical noise), normalization (to adjust for array-to-array variation), and stringent quality control to ensure data integrity.

To further validate the accuracy and consistency of the provided metadata (sample information), predicted chronological age, sex, and ethnicity were derived from the DNA methylation data using established algorithms, namely the Horvath clock [22] for age estimation, inferSex [2] for sex prediction, and inferEthnicity [2] for ethnicity prediction. This step served as an internal control to identify potential discrepancies or mislabeling within the dataset. Importantly, no additional filtering was applied based on disease status, as the primary aim was to investigate genome-wide methylation patterns without bias. Furthermore, samples from both sexes were included and analyzed together, without stratification. In subsequent analyses, sex was found to have a minimal effect compared to the methylation patterns of specific transcripts.

After all preprocessing and quality control steps, a total of 714 out of 732 samples met the eligibility criteria and were retained for subsequent analyses.

When individual CpG values were missing within a sample, those sites were imputed as zero, enabling the models to analyse the sample despite incomplete CpG coverage for that transcript. This strategy keeps each sample in the analysis even when one or more CpG sites linked to a transcript are unavailable.

### 2.4 Grouping samples by age for exploratory CpG analysis

As a supplementary analysis to provide general insights, we grouped individuals into three age categories: 14–30, 31–60, and 61–94 years. For each CpG site, mean methylation (*β*-value) was calculated within each group, and the difference between groups G3 and G1 was determined. This overview is not central to the study’s main objectives, but the results are presented for context. The full dataset is available in the supplementary files.

To generate these results, we used a MySQL query to efficiently calculate the average *β*-value for each CpG across age groups and the corresponding difference between G3 and G1. Only CpGs showing an absolute difference greater than 0.2 between these groups are included in the summary (file group_age_cpg.sql).

### 2.5 Transcript Mapping

To interpret the biological significance of age-associated methylation, we used the Illumina 450K array manifest to map CpG sites to their corresponding gene transcripts. The manifest provides detailed annotation for each probe, including genomic coordinates and transcript identifiers. Using this information, we grouped CpGs by transcript, acknowledging that a single gene may have multiple transcripts, each potentially associated with a distinct set of CpGs. Moreover, some CpGs can be annotated to more than one transcript or even to more than one gene, leading to overlaps and intercalated CpG sets.

This approach resulted in a comprehensive list in which each entry contains the gene, transcript ID, and the specific set of CpGs associated with that transcript as defined by the manifest. This transcript-centric organization provided the foundation for subsequent machine learning analysis.

### 2.6 Correlation Analysis

For each transcript’s group of CpGs, we applied four relatively fast and distinct machine learing models: glmnet (regularized linear regression), krlsRadial (non-linear kernel), brnn (neural network), and bstTree (boosted decision trees), all to model the relationship between methylation and chronological age. Transcripts whose CpG methylation profiles reached an R^2^ > 0.8 with any of these models were selected as candidates for further analysis.

For transcripts passing this threshold, we further examined each individual CpG site by computing Pearson’s and Spearman’s correlation coefficients with chronological age across all 714 samples using ugPlot. For each CpG, the squared Pearson correlation (R^2^) between methylation level (*β*-value) and age was used to quantify the strength of association. This additional step allowed us to identify CpGs that, on their own, showed strong predictive value for chronological age.

### 2.7 Model Training and Evaluation with ugPlot

To evaluate the large number of transcripts and CpGs tested in this study, we developed **ugPlot**, a custom R package designed to automate model testing using over 200 machine learning algorithms from the caret package, a widely used and well-established R package that provides a unified framework for training and evaluating predictive models. ugPlot builds on caret by offering a simplified interface through **Shiny**, allowing users to load datasets, choose target variables, and compare model performance through a graphical interface without requiring programming experience. This made it possible to test many models efficiently and reproducibly, while making the tool accessible to users beyond the bioinformatics field. Although originally developed for DNA methylation analysis, ugPlot can be applied to a wide range of structured datasets in other areas and can be downloaded from https://github.com/a00s/ugplot

Gender (presented as 0 and 1 in the dataset), though primarily used in the models, was later removed from the predictor set because runs showed that, for those specific transcripts, it was not an important feature.

If all CpGs associated with a transcript were zero in a given sample, that sample was excluded from the prediction for that transcript.

ugPlot first partitions the dataset into 80% for training and 20% for validation. Each model is then evaluated using 10-fold cross-validation to estimate its predictive performance. Because not all models in the **caret** package are compatible with every type of dataset, ugPlot attempts to fit each model and automatically skips those that fail. Some models, for instance, require exclusively numeric input, while others are designed to work with binary or categorical variables. This automated handling avoids manual filtering and ensures that only models appropriate for the given dataset are included in the final evaluation.

A key feature of **ugPlot** is the use of fixed seed values to ensure that the same analysis produces identical results every time it is run. To evaluate how different data partitions affect model performance, ugPlot was configured to run the same analysis using a sequence of 4000 seed values. From the seed that gave the best performance, we selected one and repeated the training process 4000 times to measure how much the results varied due to internal randomness. For each model, we also calculated the interquartile range (IQR) of the R^2^ values, to confirm that the observed performance was not the result of occasional outliers or random fluctuations. This use of seeds ensures that any analysis can be exactly reproduced by other researchers, as long as the same dataset and seed are used. An illustrated ugPlot workflow is shown in Figure 2.

**Figure 2.**
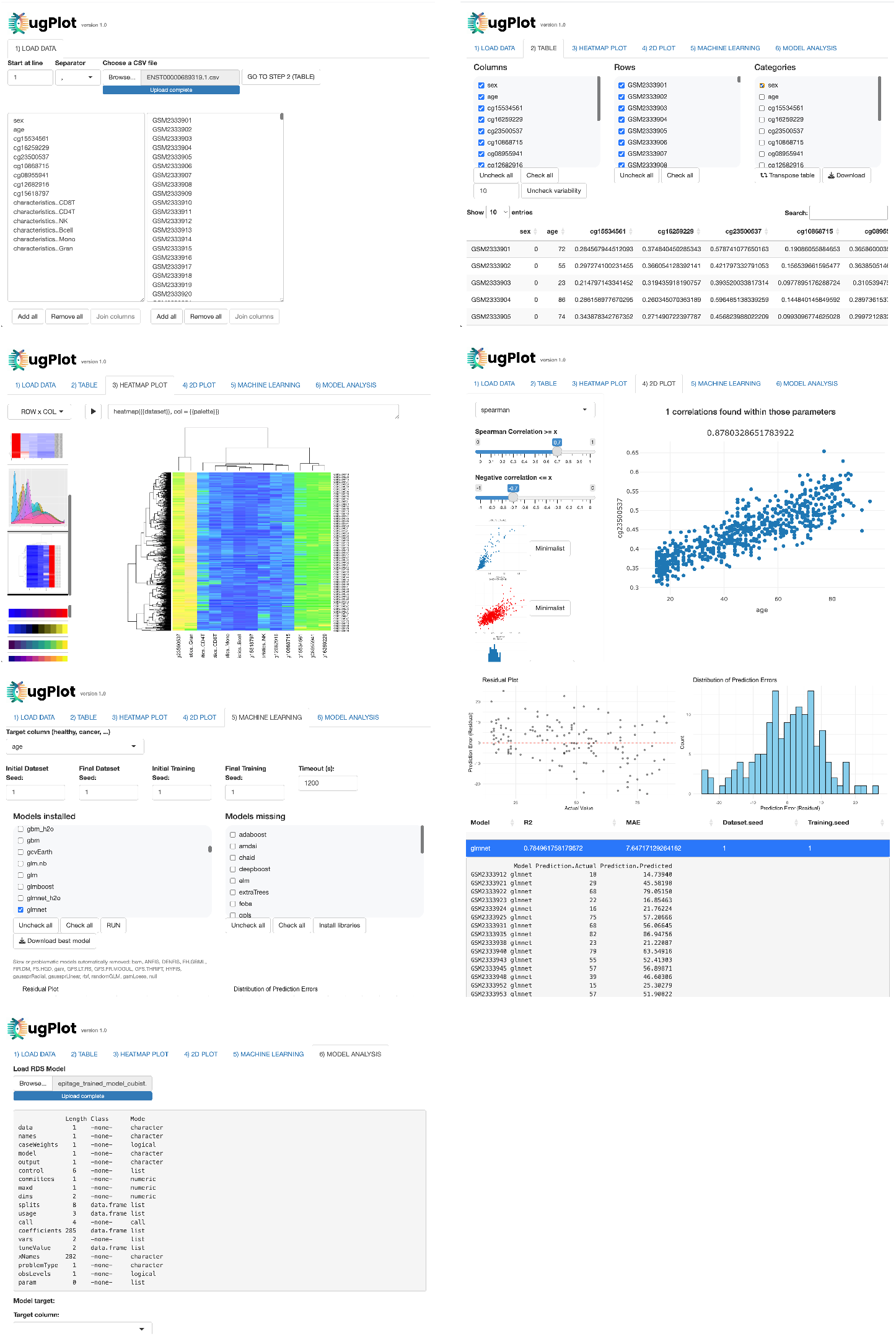
Example images illustrating the ugPlot workflow: (1) loading and merging of multiple input files; (2) dataset management and explicit definition of variable data types; (3) plot generation for exploratory analysis; (4) bidimensional visualization of CpG–age correlations; (5) automated scanning and selection of the best-performing models; (6) visualization of model performance and prediction results; and (7) application of a trained model to an external cohort.

The final output is presented in Table 2 and includes, for each transcript, the best-performing model, its corresponding seed, and the R^2^ value. Pearson’s and Spearman’s correlation coefficients are also shown when their values exceed 0.8. By automating the steps of data selection, model fitting, and result recording in a single framework, ugPlot minimizes the risk of manual errors and makes it practical to test large numbers of transcripts. Trained models were exported as.rds files and applied to an external dataset (GSE116339) to predict age, as demonstrated in this study.

## 3 Results

### 3.1 Mean CpG methylation across age groups

As an initial overview, we examined average CpG methylation levels across three age groups (14–30, 31–60, and 61–94 years). Table 1 reports the mean methylation (*β*-value) within each group (G1, G2, and G3, respectively), as well as the difference between G3 and G1, showing only CpGs with the strongest age-related differences (|Δ*β*| > 0.3). The Gene column displays the gene name annotated in the Illumina manifest; if absent, no gene annotation was available for that CpG in the manifest. Due to cohort imbalance and lack of gene-context stratification, these results should be interpreted with **caution** and are intended only as a preliminary descriptive summary. The full table is available in the supplementary files.

**Table 1.**
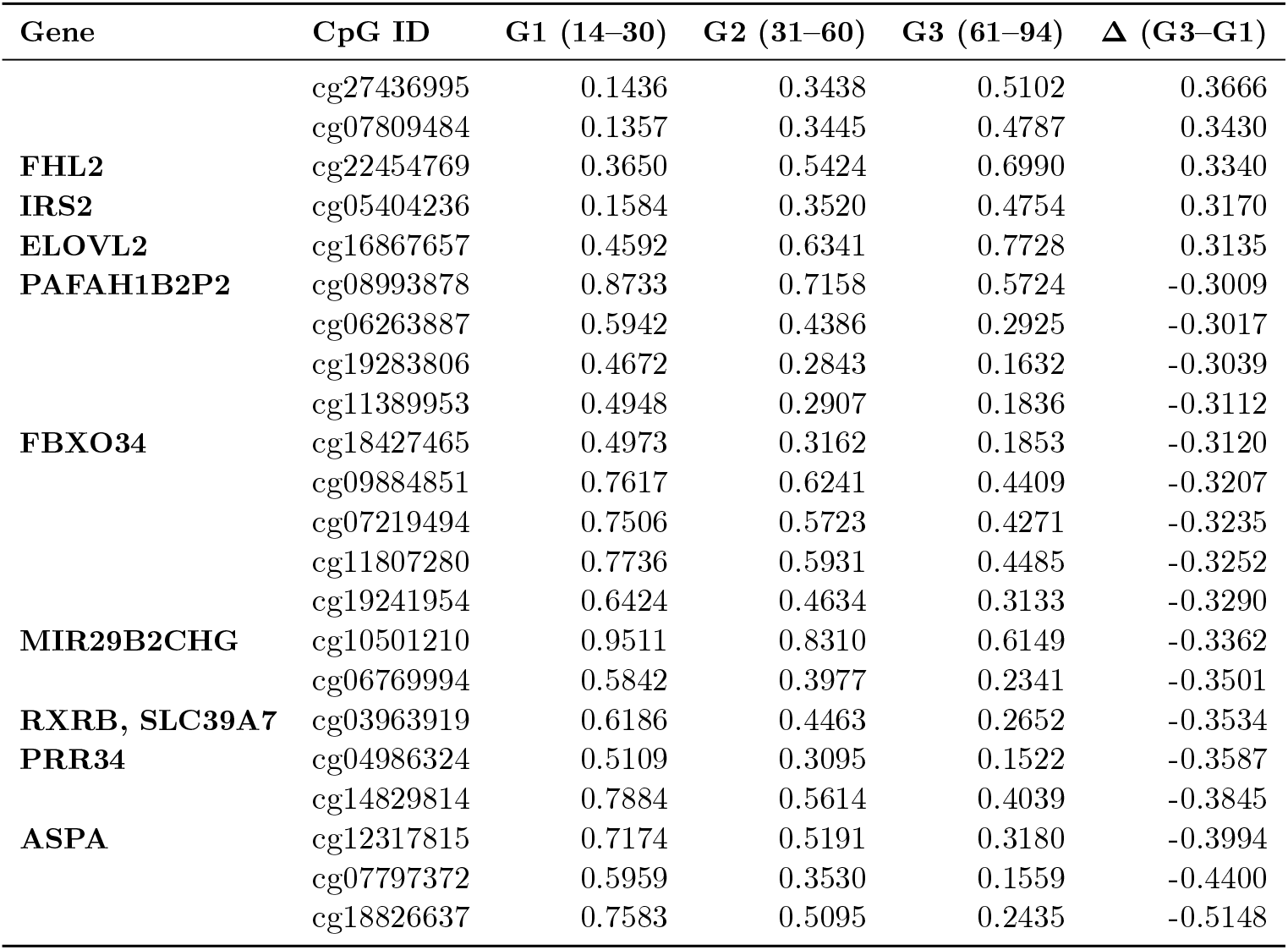
Mean CpG methylation (β-values) in three age groups, showing only CpGs with the largest age-related differences. Gene names were extracted from the Illumina manifest.

**Table 2.** Top Epitage transcripts associated with age. For each transcript, the table reports Pearson’s and Spearman’s correlations (when ≥ 0.8), NaD (percentage of absent CpGs set as zero), the best-performing machine learning model, its seed, and the range of R^2^ values across 4000 seeds, with and without inclusion of blood subpopulation proportions (wB).

| Gene | Transcript | Pearson | Spearman | Best model $R^2$ | NaD | Best model / seed / $R^2$ range |
| --- | --- | --- | --- | --- | --- | --- |
| ELOVL2 | ENST00000456616.2 | cg21572722 **0.91** | cg21572722 **0.92** | 0.95 | 0% | **bagEarthGCV** (243:1), 0.88–0.95 (IQR 0.013)<br>**wB:** (239:1), 0.87–0.95 (IQR 0.011) |
|  | ENST00000667154.1 | cg16867657 **0.95** | cg16867657 **0.95** |  |  |  |
|  | ENST00000657744.1 | cg24724428 **0.86** | cg24724428 **0.87** |  |  |  |
| ELOVL2 | ENST00000354666.4 | cg16867657 **0.95** | cg16867657 **0.95** | 0.95 | 0% | **bagEarth** (1186:1231), 0.88–0.95 (IQR 0.013)<br>**wB:** (239:1), 0.87–0.95 (IQR 0.012) |
|  |  | cg24724428 **0.86** | cg24724428 **0.87** |  |  |  |
|  |  | cg21572722 **0.91** | cg21572722 **0.92** |  |  |  |
| ELOVL2 | ENST00000667435.1 | cg16867657 **0.95** | cg16867657 **0.95** | 0.95 | 0% | **cubist** (243:3), 0.87–0.95 (IQR 0.013)<br>**wB:** (243:1), 0.90–0.95 (IQR 0.010) |
|  | ENST00000607275.6 | cg24724428 **0.86** | cg24724428 **0.87** |  |  |  |
|  |  | cg21572722 **0.91** | cg21572722 **0.92** |  |  |  |
| FHL2 | ENST00000409177.6 | cg06639320 **0.92** | cg06639320 **0.93** | 0.94 | 0% | **gamLoess** (3157:8479), 0.85–0.94 (IQR 0.019)<br>**wB:** (3157:1), 0.64–0.94 (IQR 0.023) |
|  |  | cg24079702 **0.89** | cg24079702 **0.90** |  |  |  |
|  |  | cg22454769 **0.92** | cg22454769 **0.92** |  |  |  |
| FHL2 | ENST00000393353.7 | cg24079702 **0.89** | cg24079702 **0.90** | 0.94 | 0% | **bagEarth** (2123:1849), 0.45–0.94 (IQR 0.025)<br>**wB:** (299:1), 0.54–0.94 (IQR 0.022) |
|  | ENST00000344213.9 | cg22454769 **0.92** | cg22454769 **0.92** |  |  |  |
|  | ENST00000530340.6 | cg06639320 **0.92** | cg06639320 **0.93** |  |  |  |
|  | ENST00000358129.8 |  |  |  |  |  |
|  | ENST00000408995.5 |  |  |  |  |  |
|  | ENST00000322142.13 |  |  |  |  |  |
| TRIM59 | ENST00000471155.5 | cg07553761 **0.87** | cg07553761 **0.89** | 0.90 | 0.79% | **bstTree** (2233:1), 0.74–0.90 (IQR 0.028)<br>**wB:** (2422:1), 0.75–0.89 (IQR 0.027) |
|  | ENST00000483754.1 |  |  |  |  |  |
|  | ENST00000471396.1 |  |  |  |  |  |
|  | ENST00000479460.5 |  |  |  |  |  |
|  | ENST00000309784.9 |  |  |  |  |  |
| OBSCN | ENST00000422127.5 | – | – | 0.90 | 0% | **gamboost** (3928:1), 0.73–0.90 (IQR 0.029)<br>**wB:** (3763:1), 0.77–0.91 (IQR 0.025) |
|  | ENST00000680850.1 |  |  |  |  |  |
|  | ENST00000570156.7 |  |  |  |  |  |
| MIR29B2CHG | ENST00000655169.1 | – | cg10501210 **-0.86**<br>(negative) | 0.90 | 0.54% | **ranger** (3247:3210), 0.69–0.90 (IQR 0.032)<br>**wB:** (3247:1), 0.77–0.90 (IQR 0.025) |
| TRIM59 | ENST00000543469.1 | cg07553761 **0.87** | cg07553761 **0.89** | 0.88 | 5.47% | **rf** (2233:1), 0.74–0.88 (IQR 0.029)<br>**wB:** (1417:1), 0.75–0.89 (IQR 0.028) |
| TRIM59 | ENST00000494486.1 | cg07553761 **0.87** | cg07553761 **0.89** | 0.88 | 0.78% | **bagEarth** (3419:1815), 0.43–0.88 (IQR 0.040)<br>**wB:** (3419:1), 0.48–0.90 (IQR 0.034) |
| TRIM59 | ENST00000496222.1 | cg07553761 **0.87** | cg07553761 **0.89** | 0.88 | 0.79% | **bagEarth** (1870:1), 0.43–0.88 (IQR 0.043)<br>**wB:** (3419:1), 0.39–0.90 (IQR 0.036) |
| MIR29B2CHG | ENST00000487977.2<br>ENST00000652846.1<br>ENST00000702741.1 | – | cg10501210 **-0.86** | 0.89 | 0.75% | **ranger** (3247:2470), 0.72–0.89 (IQR 0.032)<br>**wB:** (3247:1), 0.79–0.90 (IQR 0.025) |
| OBSCN | ENST00000636476.2 | – | – | 0.89 | 0% | **gamboost** (3928:1), 0.68–0.89 (IQR 0.036)<br>**wB:** (3526:1), 0.76–0.89 (IQR 0.029) |
| KLF14 | ENST00000583337.4 | cg08097417 **0.80** | cg08097417 **0.83**<br>cg07955995 **0.83** | 0.88 | 10.31% | **ppr** (1699:1), 0.69–0.88 (IQR 0.035)<br>**wB:** (1734:1), 0.76–0.92 (IQR 0.029) |
| PRRT1 | ENST00000472641.1 | – | – | 0.88 | 5.70% | **brnn** (548:4), 0.65–0.88 (IQR 0.043)<br>**wB:** (2824:1), 0.67–0.89 (IQR 0.036) |
| PRRT1 | ENST00000375150.6 | – | – | 0.88 | 1.75% | **svmLinear3** (1181:1), 0.66–0.88 (IQR 0.041)<br>**wB:** (1181:1), 0.72–0.89 (IQR 0.033) |
| OBSCN | ENST00000662438.1 | – | – | 0.88 | 0% | **ppr** (3526:1), 0.64–0.88 (IQR 0.041)<br>**wB:** (2943:1), 0.73–0.90 (IQR 0.030) |
| CALB1 | ENST00000518457.5 | cg26290632 **0.86** | cg26290632 **0.88** | 0.87 | 0.82% | **ppr** (3526:1), 0.69–0.87 (IQR 0.037)<br>**wB:** (329:1), 0.77–0.91 (IQR 0.024) |
| CALB1 | ENST00000476853.1<br>ENST00000473670.1 | cg26290632 **0.86** | cg26290632 **0.88** | 0.87 | 0.07% | **bagEarth** (2192:1), 0.22–0.87 (IQR 0.039)<br>**wB:** (1727:1), 0.55–0.89 (IQR 0.030) |
| CALB1 | ENST00000265431.7 | cg26290632 **0.86** | cg26290632 **0.88** | 0.87 | 1.20% | **bagEarth** (2192:1), 0.45–0.87 (IQR 0.042)<br>**wB:** (329:1), 0.49–0.89 (IQR 0.032) |
| CALB1 | ENST00000482702.5 | cg26290632 **0.86** | cg26290632 **0.88** | 0.87 | 0.08% | **bagEarth** (2192:1), 0.35–0.87 (IQR 0.040)<br>**wB:** (1727:1), 0.60–0.90 (IQR 0.030) |
| OBSCN | ENST00000284548.16 | – | – | 0.87 | 0% | **brnn** (930:1), 0.59–0.87 (IQR 0.038)<br>**wB:** (3625:1), 0.66–0.88 (IQR 0.034) |
| OTUD7A | ENST00000678495.1 | cg04875128 **0.84** | cg04875128 **0.84**<br>cg01763090 **0.81** | 0.87 | 0% | **bagEarth** (645:487), 0.66–0.86 (IQR 0.044)<br>**wB:** (104:1), 0.72–0.89 (IQR 0.034) |
| PRRT1 | ENST00000211413.10 | – | – | 0.87 | 4.66% | **glmnet** (1215:1), 0.61–0.87 (IQR 0.037)<br>**wB:** (1181:1), 0.67–0.88 (IQR 0.032) |
| PRRT1 | ENST00000495191.5 | – | – | 0.87 | 4.52% | **glmboost** (977:3), 0.65–0.87 (IQR 0.033)<br>**wB:** (2824:1), 0.69–0.88 (IQR 0.029) |
| KCNS1 | ENST00000537075.3 | cg19702785 **0.78** | cg19702785 **0.80** | 0.87 | 0.01% | **ppr** (938:1), 0.65–0.87 (IQR 0.042)<br>**wB:** (247:1), 0.73–0.89 (IQR 0.035) |
| VTRNA1-2 | ENST00000689319.1 | cg23500537 **0.86** | cg23500537 **0.87** | 0.86 | 2.58% | **bstTree** (577:6), 0.68–0.86 (IQR 0.035)<br>**wB:** (577:1), 0.69–0.87 (IQR 0.033) |
| OTUD7A | ENST00000560598.2 | cg04875128 **0.84** | cg04875128 **0.84**<br>cg01763090 **0.81** | 0.87 | 0% | **bstTree** (130:1), 0.65–0.87 (IQR 0.042)<br>**wB:** (3173:1), 0.67–0.89 (IQR 0.038) |
| PRRT1 | ENST00000467780.5 | – | – | 0.86 | 0.15% | **brnn** (1578:1), 0.66–0.86 (IQR 0.036)<br>**wB:** (627:1), 0.74–0.90 (IQR 0.026) |
| SYNGR3 | ENST00000248121.7 | cg11220950 **0.80** | cg11220950 **0.81** | 0.85 | 0% | **cubist** (268:45), 0.63–0.85 (IQR 0.046)<br>**wB:** (3488:1), 0.67–0.88 (IQR 0.031) |
| SPTBN4 | ENST00000352632.7 | – | – | 0.82 | 0.11% | **gamboost** (2155:1), 0.58–0.82 (IQR 0.042)<br>**wB:** (2155:1), 0.64–0.85 (IQR 0.036) |

### 3.2 Top Transcripts Associated with Ageing

Applying correlation and model performance cutoffs, we found 48 gene transcripts significantly correlated with age. These associations were determined either by the presence of at least one CpG with Pearson’s or Spearman’s correlation coefficient (|r| ≥ 0.8), or by overall transcript-level model performance exceeding R^2^ = 0.8. Table 2 presents a summary of these top transcripts along with their statistical measures. Our results include several genes already linked to ageing, as well as others with limited prior evidence or indirect associations. This approach ensures that only the strongest associations identified by each method are reported.

As the 450K array annotates some CpGs to multiple transcript isoforms, identical CpG sets may appear under different transcript IDs, resulting in shared statistics for these entries. This can occur when two transcripts are originally annotated with mostly, but not entirely, the same CpGs, and the CpG(s) that would make one transcript unique lack data across all samples. In such cases, both transcripts effectively share the same CpG set with available information and are treated identically in the analysis.

The genes ELOVL2 [15] and FHL2 [24] consistently ranked among the top transcripts identified, aligning with their established roles as benchmark markers in epigenetic ageing studies. In contrast, KCNS1, SPTBN4, and VTRNA1-2; genes rarely cited in the methylation-ageing literature; demonstrated R^2^ values comparable to those well-established markers, highlighting their potential as novel candidates for future investigation. The analysis also revealed instances of transcript redundancy; for example, several ELOVL2 isoforms share the same three CpGs (cg16867657, cg21572722, cg24724428). The optimal models identified encompassed a range of algorithmic families, including linear (glmnet), nonlinear (gamboost, bagEarthGCV), neural network (brnn), and tree-based (bstTree, cubist) methods, further emphasizes the advantage of screening a broad range of machine learning approaches rather than relying on a single method. Importantly, the top-performing transcripts exhibited robust results, with interquartile ranges (IQR) of R^2^ values across 4000 resampling runs consistently below 0.04, suggesting that the observed associations are not artifacts of outliers or random data splits.

Most of the results are consistent with previous studies on established ageing genes, pointed out in the “Discussion session” of this paper, while a subset of transcripts with strong associations in our analysis have not been reported, to our knowledge, in the existing literature. This combination underscores both the robustness of the approach and the potential to identify new targets related to epigenetic ageing.

In the Table 2, R^2^ values for the best models were obtained using the postResample() function from the caret R package, where R^2^ is calculated as “the square of the correlation between the observed and predicted outcomes”, as documented in the official manual [26]. Corresponding plots for each transcript are presented in Figure 3.

**Figure 3.**
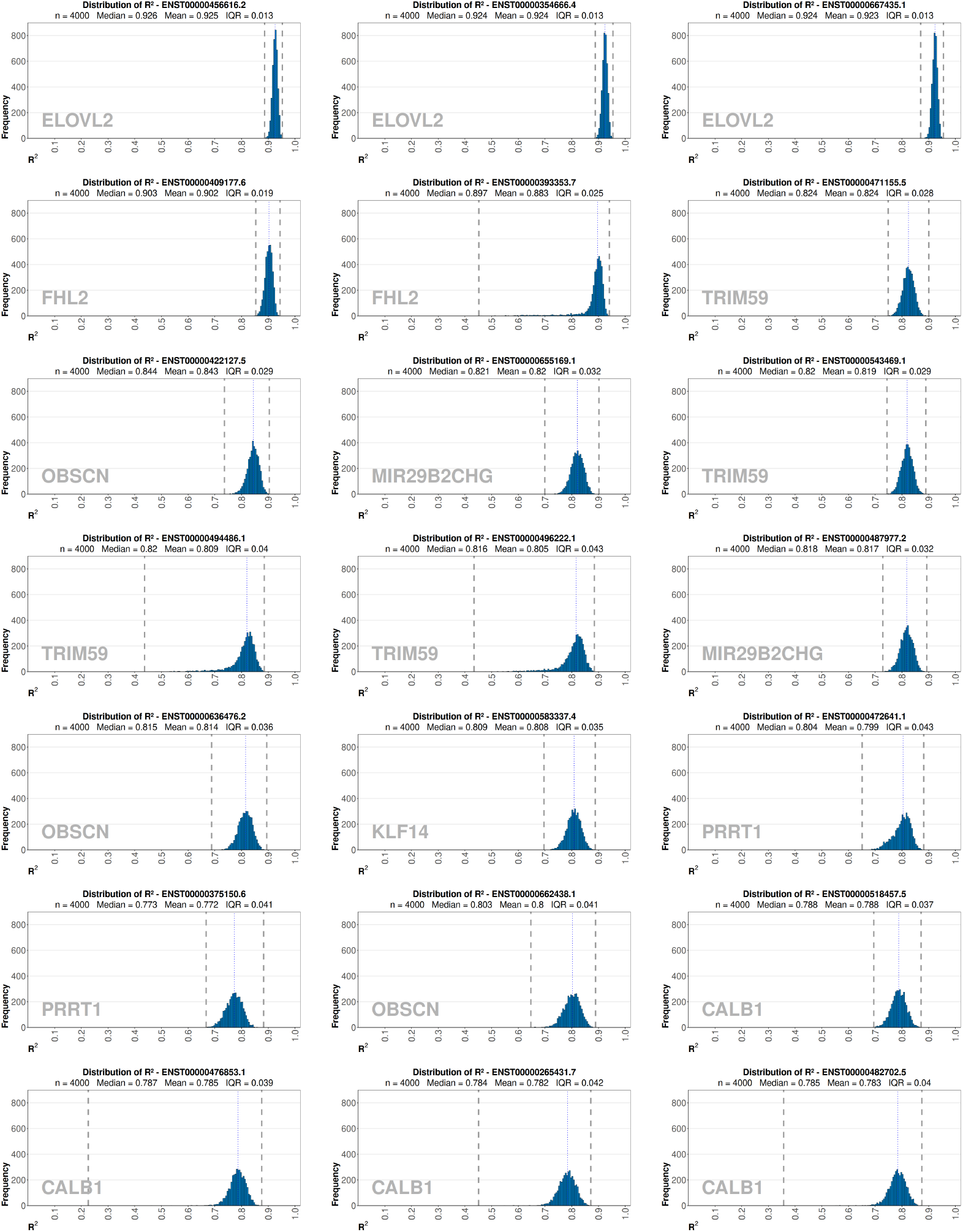
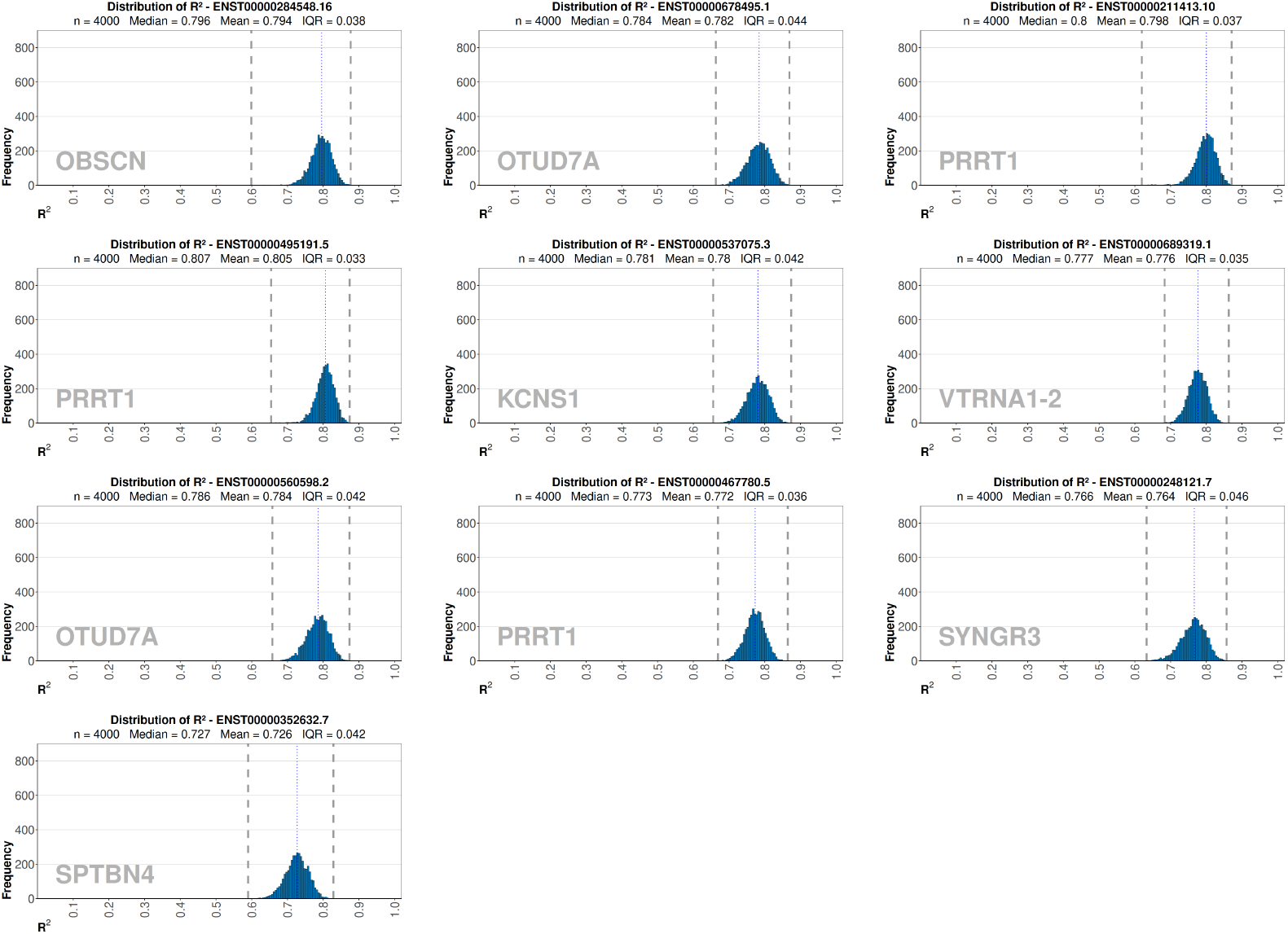
Distribution of R^2^ values for each Epitage transcript, obtained from 4000 sequential seeds for train/test splits. Each histogram illustrates the robustness of the predictive model in relation to dataset sample size. The dashed lines indicate the minimum and maximum R^2^ values observed. Plots were generated using the ugPlot R package.

### 3.3 Consequences of zero imputation for missing CpG values

Among the 714 usable samples, some CpGs were absent or failed quality control, resulting in missing values. Zero imputation was adopted for computational efficiency and to avoid the introduction of categorical indicators, which would reduce the number of machine learning models available for analysis.

To evaluate whether zero imputation influenced model performance, we repeated the analysis using datasets where any CpG with at least one missing value was removed across all samples. The only exception was the gene MIR29B2CHG, which shows a very promising CpG that was missing in 27 samples, making it more reasonable to remove the samples instead of the CpG..

As expected, removing CpGs with missing values across all samples led in some cases to reduced performance or increased IQR, as informative CpGs were excluded even from samples with otherwise valid measurements. Nevertheless, under this more stringent approach, all transcripts continued to meet the R^2^ ≥ 0.8 selection criterion, suggesting that the models naturally assigned lower weight to CpGs with a high proportion of missing values.

### 3.4 Blood subpopulations and age prediction

The composition of major blood cell subpopulations undergoes significant changes throughout the human lifespan, reflecting processes such as immunosenescence [36]. Age-related remodeling of the immune system typically involves shifts in the proportions of CD4+ and CD8+ T cells, B cells, NK cells, monocytes, and neutrophils [36]. Recent studies using DNA methylation and flow cytometry have confirmed that both cell type proportions and intrinsic cell properties change with ageing, and that these main subpopulations, including monocytes, B cells, and T cell subsets, exhibit distinct biological ages and methylation profiles, which can impact the interpretation of epigenetic clocks measured in whole blood [33]; [36].

This next analysis was performed to rule out the possibility that transcript-based models were not truly predicting age, but instead were primarily capturing information about blood cell subpopulation composition, which is itself correlated with age. To test this, we removed all CpGs from the dataset and built models using only the proportions of six major leukocyte types, CD8+ T cells, CD4+ T cells, natural killer (NK) cells, B cells, monocytes, and granulocytes, from the GSE87571 dataset. If blood cell composition alone explained age prediction, these models would be expected to perform as well as transcript-based models. However, the performance of the subpopulation-based models was lower, with a maximum R^2^ from 0.29 to 0.62 across 4000 random train/test splits with IQR of 0.066 using bagEarth model (Figure 4). Still, the predictive power of blood cell composition is not negligible, meaning that even though transcript-based models perform better, part of their predictive accuracy may still depend on differences in cell subpopulations.

**Figure 4.**
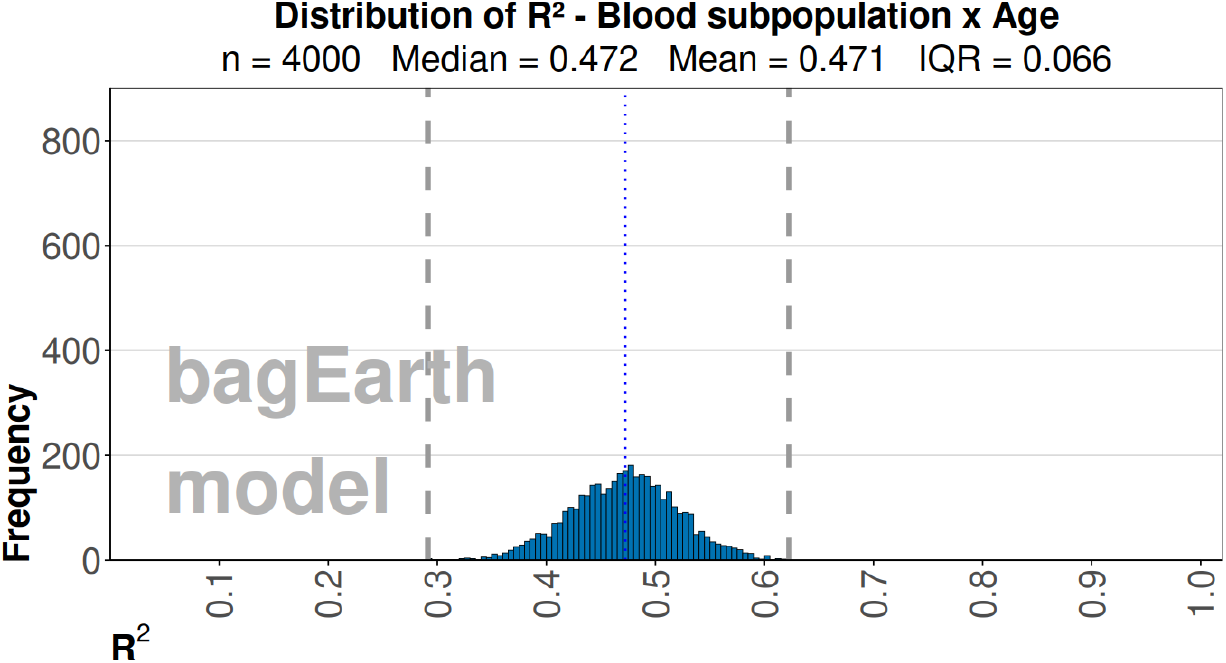
Distribution of R^2^ from 4000 seeds predicting age with blood cell subpopulations using the bagEarth model. Dashed lines show minimum and maximum values. Plot generated with ugPlot.

Furthermore, adding blood cell subpopulation variables to transcript-based methylation models led to an improvement in R^2^ of up to 0.03 (on a scale where the maximum possible value is 1). Among these variables, the NK cell proportion often emerged as particularly relevant, sometimes reaching an importance value greater than 0.5 when looking at ugPlot reports for which parameters were most relevant in the result. This suggests that, while blood cell subpopulations alone are not strong predictors of age, they can enhance model performance when combined with transcript data.

We then investigated whether this effect could be due to the direct predictive value of the abundance of specific cell types in relation to ageing, or because these variables help the model contextualize CpG signals based on cell composition. For each transcript, we included its CpGs together with one cell subtype at a time, excluding sex and age information, to test whether the CpGs alone could predict the cell subpopulation. We repeated this process for all six cell subtypes from the dataset and for every transcript on the Epitage list. The maximum R^2^ obtained barely reached 0.61, showing again that when looking at transcripts alone, the relationship with cell subpopulations, although present, is not large enough to explain the age prediction through cell composition.

Then we tried to predict the cell subpopulation by merging all 331 CpGs into a single dataset without age and sex. The best result was obtained for granulocytes using the enet model, reaching a maximum R^2^ of 0.84 (Table 3) with the respective plots shown in Figure 5. Most of the result was influenced by multiple CpGs, mainly from the genes PRRT1 and OBSCN. This shows that, although some genes, in combination, can predict cell subpopulations to some extent, it is unlikely that cell subpopulations individually influenced the result for the transcript list used in Epitage.

**Table 3.** Performance of predictive models for the blood cell subpopulations available in the dataset, obtained using the 331 CpGs from the Epitage list simultaneously and evaluated across 4000 sequential seeds for train/test splits.

| Cell type | Best model | $R^2$ range | IQR | Dataset seed |
| --- | --- | --- | --- | --- |
| CD8T | svmPoly | 0.29–0.69 | 0.078 | 2186 |
| CD4T | cubist | 0.35–0.73 | 0.064 | 3435 |
| NK | cubist | 0.43–0.81 | 0.057 | 3705 |
| B cell | cubist | 0.13–0.63 | 0.097 | 676 |
| Mono | glmnet | 0.00–0.30 | 0.062 | 2007 |
| Gran | enet | 0.50–0.84 | 0.049 | 1565 |

**Figure 5.**
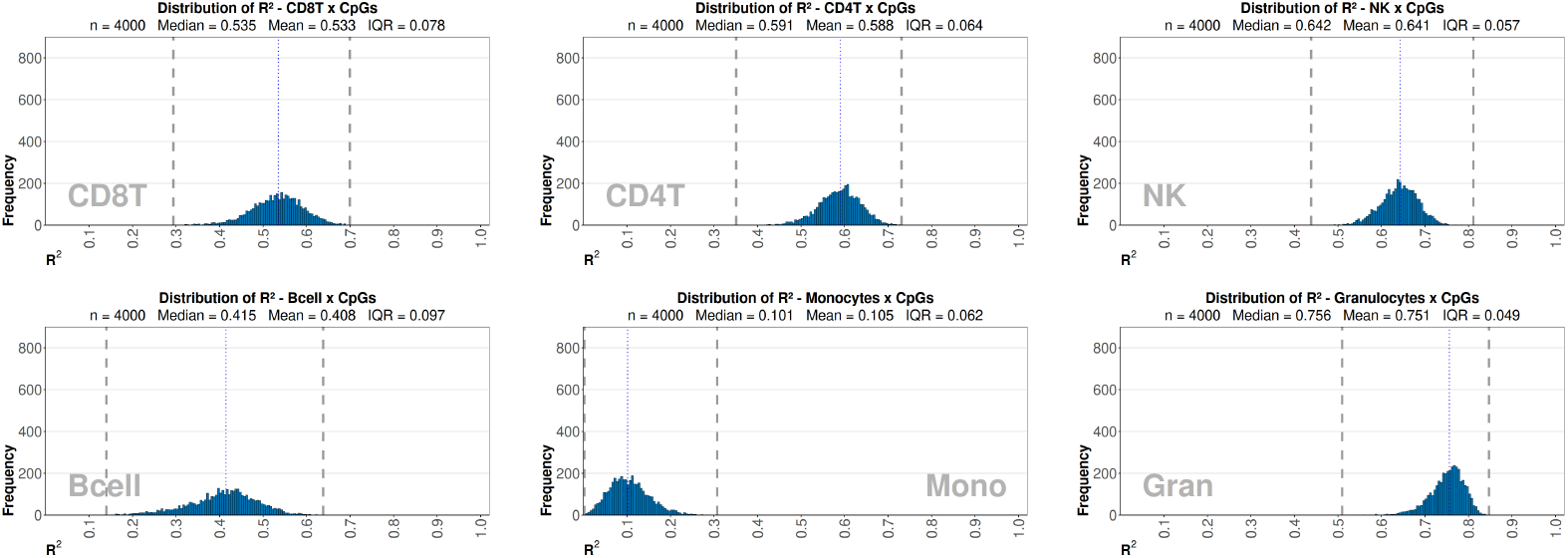
Histograms showing the distribution of R^2^ values across 4000 sequential seeds for train/test splits when predicting blood cell subpopulations (CD8T, CD4T, NK, B cells, monocytes, granulocytes) using the 331 CpGs from the Epitage list. Dashed lines indicate the minimum and maximum values obtained. Plots were generated with the ugPlot R package.

### 3.5 Validation of ugPlot transcript models for age prediction using an external cohort

To assess the robustness of the transcript-based models developed with ugPlot, each trained model for each transcript (saved as.rds files) was applied to matched CpG data from the independent dataset GSE116339 “Exposure to polybrominated biphenyl (PBB) associates with genome-wide DNA methylation differences in peripheral blood,” [10], 679 samples from 659 individuals, aged 23–88 years. The results are reported in Table 4, with corresponding plots in Figure 6.

**Table 4.**
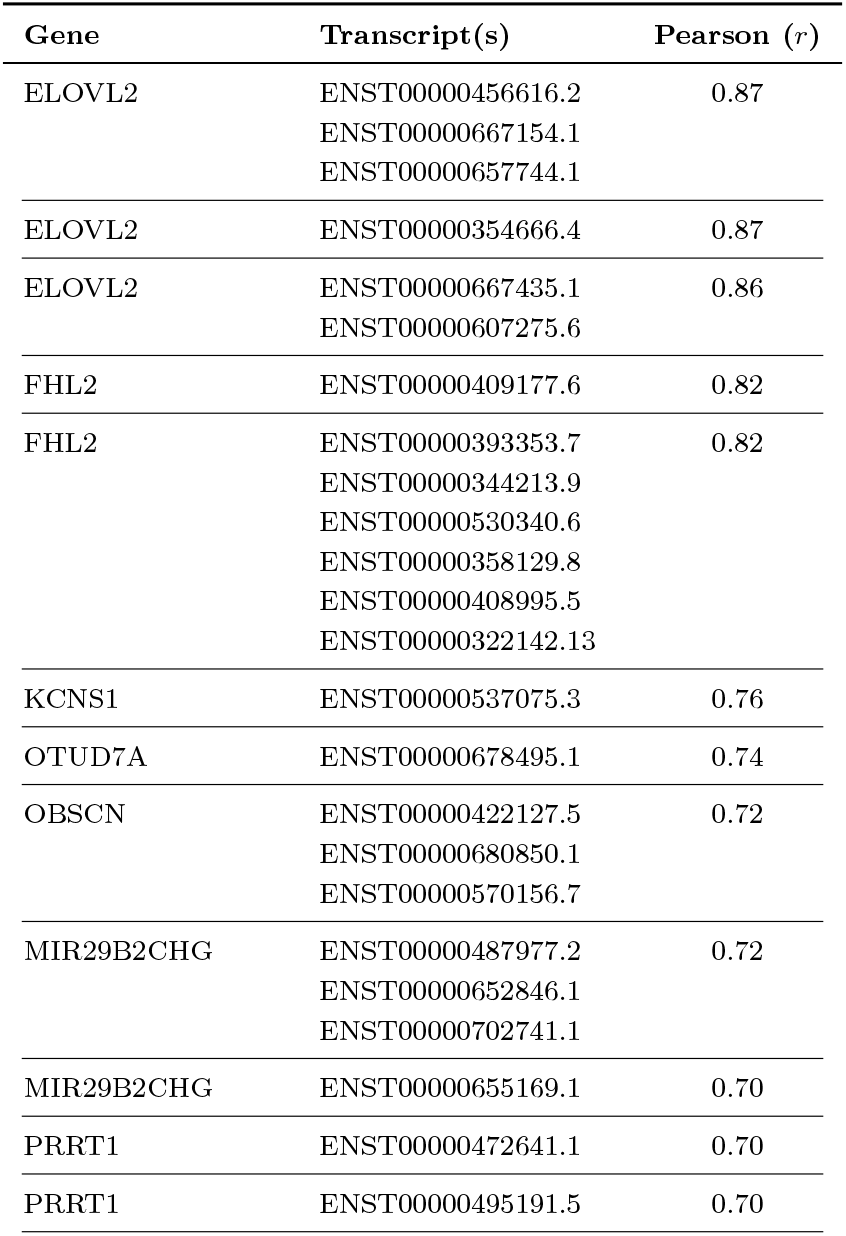
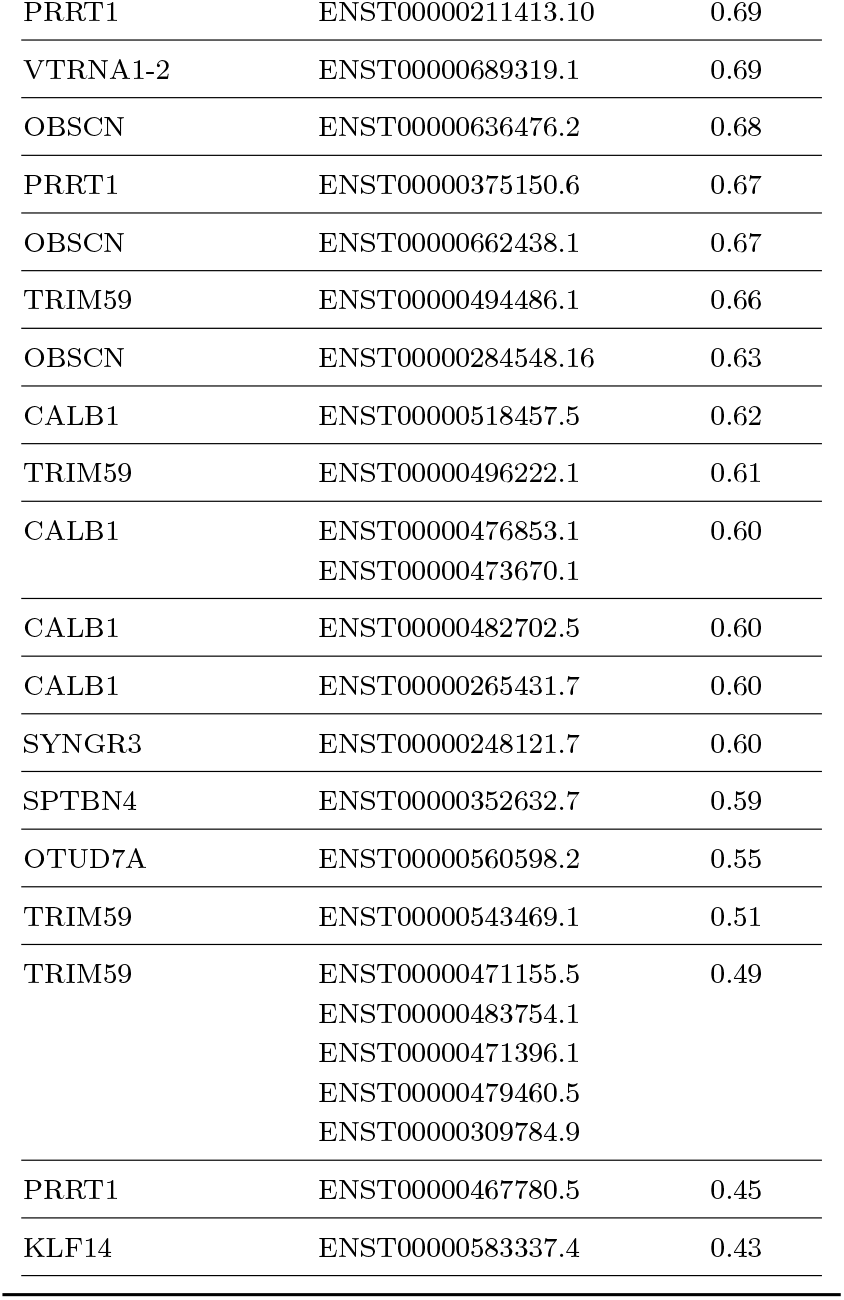
Pearson correlations between chronological age and predictions obtained by applying ugPlot pre-trained transcript models from the Epitage list to an independent external cohort GSE116339.

**Figure 6.**
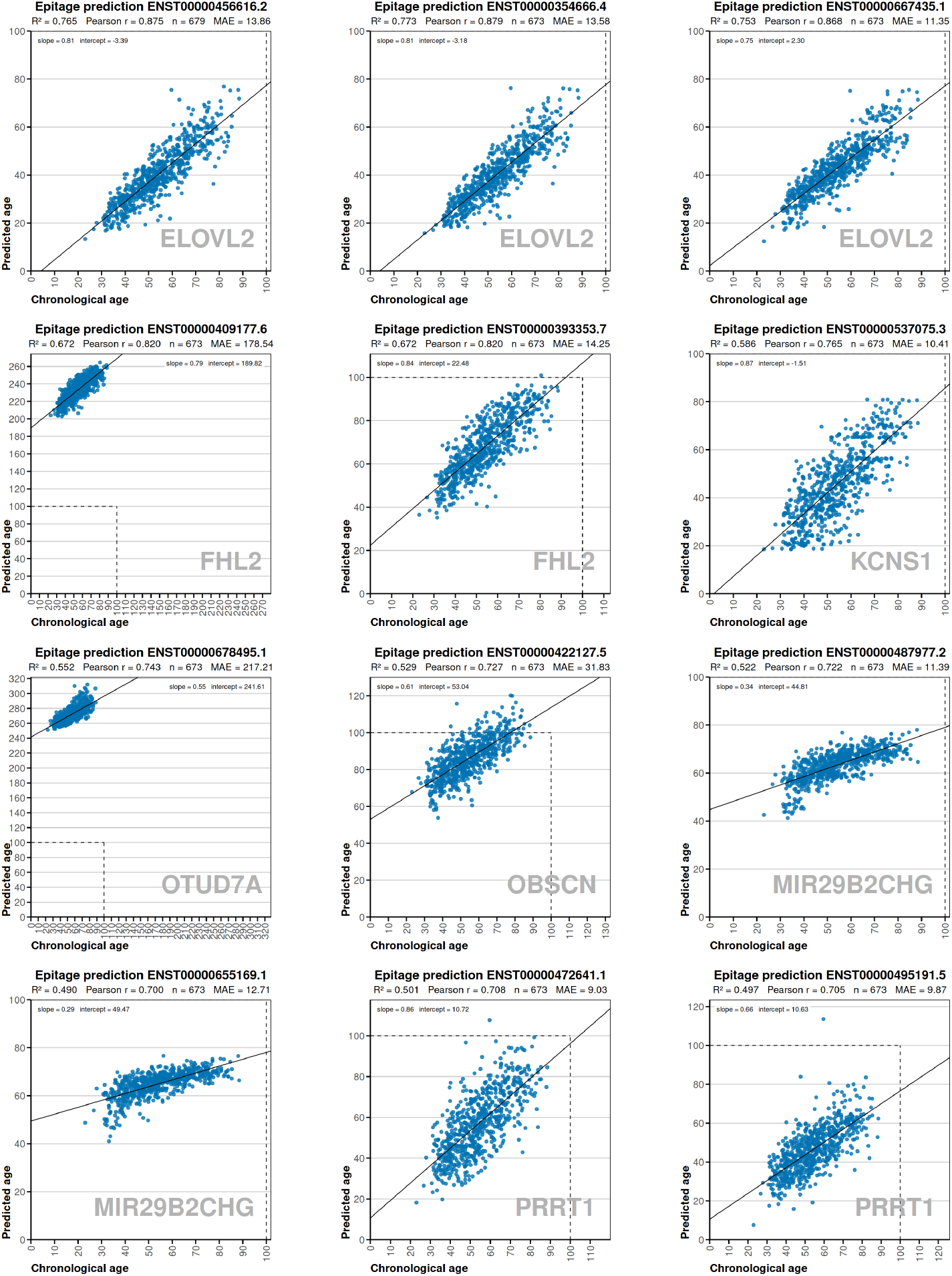
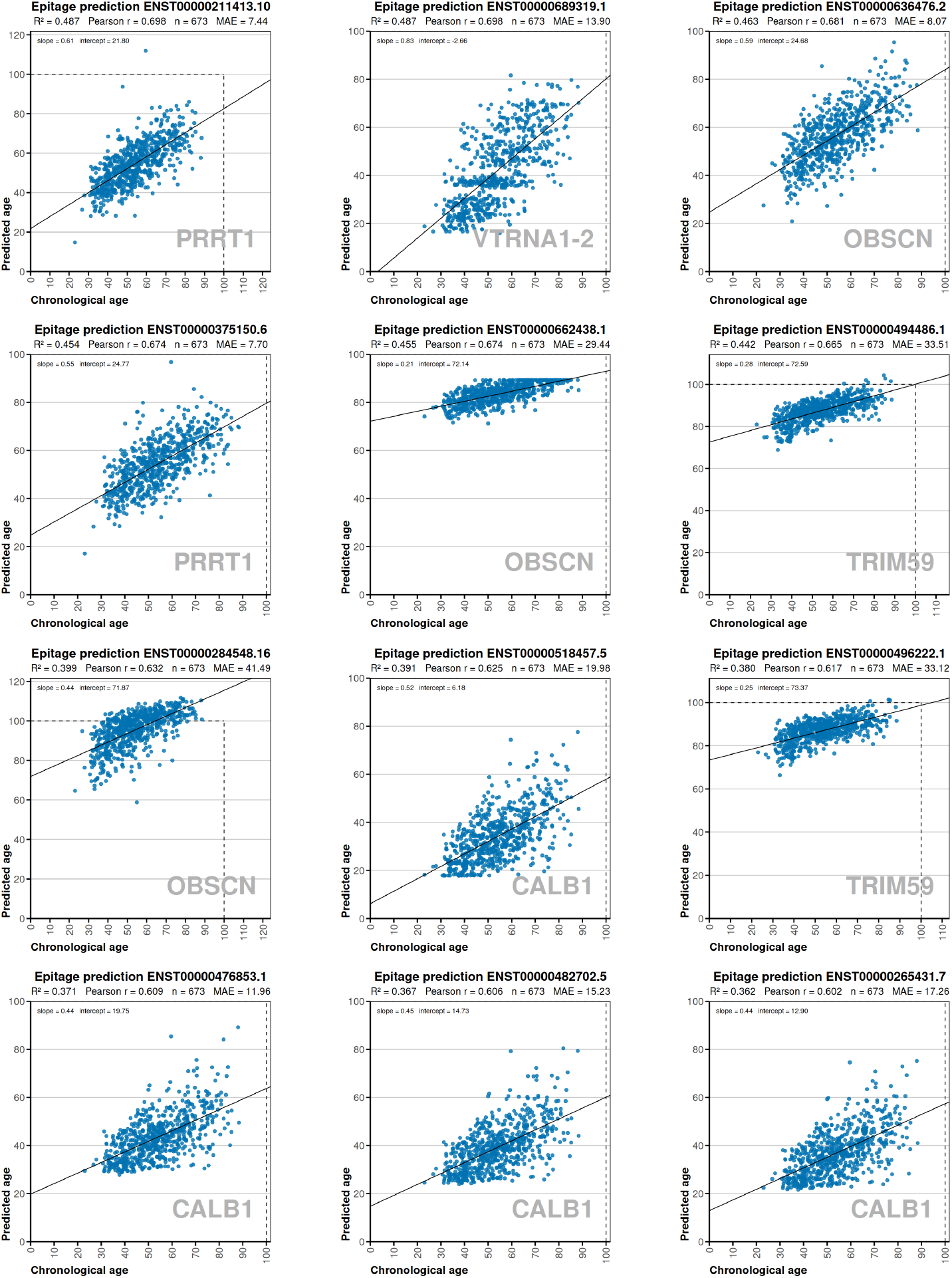
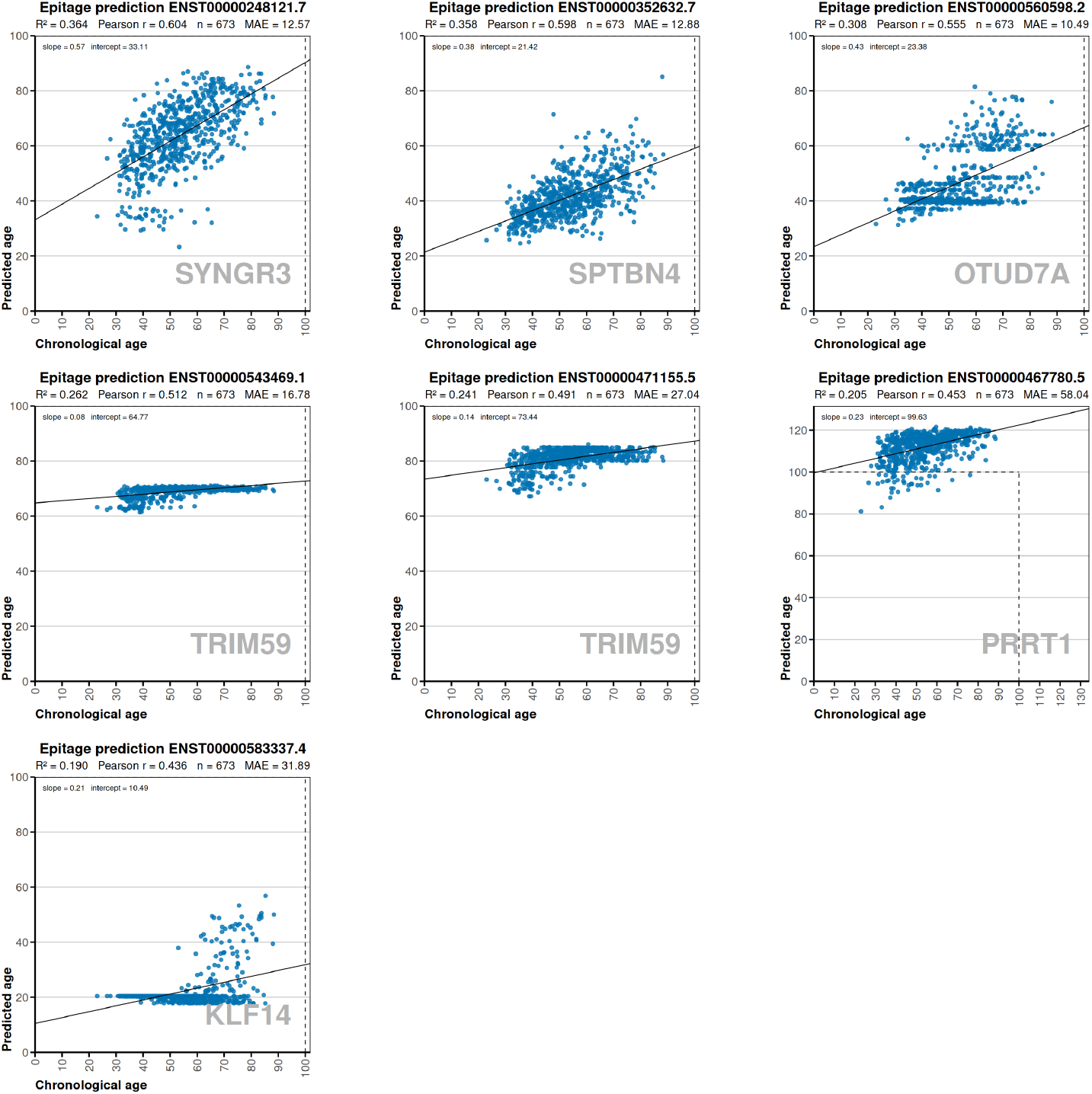
Scatter plots of predicted versus chronological age for selected Epitage transcripts in the external validation cohort. A dashed line indicates the 100-year square limit, added to highlight deviations when the plot axis was extended beyond 100. R^2^ represents the fit of the linear regression trendline. Plots were generated using the ugPlot R package.

While this dataset’s large sample size and detailed age metadata support its use for validation, prior reports indicate that PBB exposure can alter methylation and may influence age predictions as seen in “Environmental exposure to polybrominated biphenyl (PBB) associates with an increased rate of biological aging” [11].

The primary goal of this transcript-based approach is to identify transcripts with robust associations to ageing, rather than to achieve precise age prediction, which is already accomplished by established epigenetic clocks [22]. For each transcript, predictions were generated when at least one relevant CpG value was available in a given sample; samples lacking CpG data for that transcript were excluded. Application of the ugPlot-trained models to the GSE116339 dataset showed positive correlations between predicted and chronological age for most transcripts. Well-known ageing markers such as ELOVL2 and FHL2 maintained high correlations, while novel candidates like KCNS1 and SPTBN4 also showed some association with age in this independent cohort. Incomplete CpG coverage for some transcripts may have limited prediction accuracy, suggesting that future studies with broader methylation data could further refine these findings.

## 4 Discussion

There is evidence that alternative splicing produces transcript variants with distinct or even opposing functions. One example is the gene HSF4 [42], which produces both an activator and a repressor of heat shock genes through alternative splicing. Although *HSF4* did not rank among the top transcripts in the human Epitage list, it has been identified as a significant epigenetic marker in mouse ageing clocks [18]. This shows that focusing exclusively on promoter regions may overlook important regulatory signals in other parts of the gene, especially when transcripts have divergent functions.

From the literature, several genes show robust and reproducible associations with ageing at the epigenetic or expression level across diverse tissues. **ELOVL2, FHL2, KLF14, TRIM59**, and **MIR29B2CHG** consistently display strong age-related DNA methylation changes in blood with ELOVL2, KLF14, and TRIM59 showing the most consistent correlations across tissues [24]. **PRRT1** expression declines with age in both human Alzheimer’s blood and mouse hippocampus [30]; [12]. **OTUD7A** exhibits robust age-associated hypermethylation in blood, confirmed in multiple EWAS meta-analyses [44]; [3]. **SPTBN4** is subject to promoter hypermethylation and reduced expression with Alzheimer’s disease in mouse and human cortex [38]; [47]. **SYNGR3** shows an age-associated decrease in expression in both human and mouse brain [21]. **CALB1** expression is directly correlated with improved recognition memory performance in mice, supporting a role in maintaining cognitive function during ageing [40], while its age-related decrease in the rat eye lens may contribute to cataract development [16]. Copy number variants of **OBSCN** are more frequently observed in young individuals compared to older adults, suggesting a potential role for this gene in determining lifespan [17]. **vtRNA1-2** expression is influenced by epigenetic modifications, as promoter methylation has been shown to inversely correlate with its expression levels [6]. Finally, **KCNS1** is among the genes with age-associated hypermethylation across multiple immune cell types [37].

An additional aspect that deserves emphasis is the general philosophy behind ugPlot: the systematic testing of a wide range of machine learning models. In epigenetic studies, it is sometimes unclear a priori which modeling approach will perform best, as the optimal choice may depend on the dimensionality of the dataset, the correlation structure among CpGs, or the presence of non-linear effects. By automating the comparison of dozens of candidate models, ugPlot reduces the risk of bias introduced by the researcher’s prior assumptions or by reliance on a single favored algorithm. This strategy also makes it possible to detect cases where non-linear or ensemble-based methods capture hidden patterns that would otherwise remain invisible to linear models. ugPlot transforms model selection from a manual and error-prone process into a reproducible pipeline. Instead of investing substantial effort in fine-tuning one model at a time, researchers can obtain an overview of which families of algorithms are most promising for their dataset, and then focus their attention on refining the best-performing candidates.

## 5 Limitations of the Present Study

This study has some limitations that should be considered when interpreting the results. First, all analyses were based on a single whole-blood dataset (GSE87571), and methylation changes with age can be tissue-specific. Second, the Illumina 450K array, although widely used, does not cover all CpGs in the genome, and some transcripts may be underrepresented or missing due to probe coverage gaps. Additionally, the correlation-based approach identifies associations but does not establish causation or address whether methylation changes lead to altered gene expression or ageing phenotypes. Finally, although external validation was performed, differences in environmental exposures, cohort characteristics, and incomplete CpG coverage in the validation dataset may have reduced model performance. Future studies using multiple tissues, more comprehensive methylation profiling, and direct cell-type quantification will be important to confirm and extend these findings.

## 6 Conclusion

In this study, we systematically analyzed transcript-level DNA methylation profiles in human blood and identified 48 gene transcripts whose methylation is strongly associated with chronological age. By grouping CpGs by transcript and applying a broad set of machine learning models, we confirmed known ageing markers (such as **ELOVL2** and **FHL2**) and also found potential new candidates, including **KCNS1. SPTBN4**, and **vtRNA1-2**.

To facilitate large-scale and reproducible analysis, we developed **ugPlot**, an open-source R package that automates the training and evaluation of machine learning models for methylation data. The use of ugPlot enabled efficient candidate selection and ensured consistent model evaluation across multiple runs and datasets.

In summary, transcript-level methylation analysis enables the identification of gene transcripts with potential relevance to ageing. The Epitage list highlights transcripts that may be involved in ageing processes, either as active contributors or passive markers. Recognizing these transcripts not only helps to clarify the molecular pathways underlying human ageing but also opens possibilities for experimental interventions, such as restoring the expression of proteins that are no longer produced in aged tissues.

## Acknowledgments

We thank Francesco Reggiani for insightful discussions and guidance throughout the development of the project, which were essential for it to reach its current stage, and Vincenzo Fontana for guidance in the initial stages of the research. We also express our gratitude to the University of Genoa (UNIGE), Italy, for providing the connections and institutional environment that made this work possible.

## Declarations

## Ethics and consent to participate

Not applicable. This study exclusively used publicly available, anonymized datasets obtained from the Gene Expression Omnibus (GEO).

## Availability of data and materials

All datasets analyzed during this study are publicly available from the Gene Expression Omnibus (GEO) under accession numbers GSE87571 and GSE116339. The ugPlot source code used in this study is available at https://github.com/a00s/ugplot.

## Authors’ contributions

Both authors contributed substantially to this work. TBM contributed to the execution of the study, while UP provided scientific supervision, expert guidance, and critical revision of the manuscript.

## Declaration of generative AI and AI-assisted technologies in the writing process

During the preparation of this manuscript, ChatGPT was used cooperatively to improve the linguistic clarity and readability of the text. All research design, data processing, analysis, and scientific interpretation were conducted exclusively by the authors, with the support of ugPlot (a custom tool developed in the context of this study).

